# poreTally: run and publish *de novo* Nanopore assembler benchmarks

**DOI:** 10.1101/424184

**Authors:** Carlos de Lannoy, Judith Risse, Dick de Ridder

**Affiliations:** Bioinformatics Group, Wageningen University, 6700AP, Wageningen, The Netherlands; Netherlands Institute of Ecology (NIOO-KNAW), 6708PB, Wageningen, The Netherlands

## Abstract

Nanopore sequencing is a novel approach to nucleic acid analysis that generates long, error-prone reads. Since device components, base calling software and best practices for sample preparation are updated frequently and extensively, the nature of the produced data also changes frequently. As a result, peer-reviewed publications on *de novo* assembly pipeline benchmarking efforts are quickly rendered outdated by the next major improvement to the sequencing platforms. To provide the user community with a faster, more flexible alternative to peer-reviewed benchmark papers for *de novo* assembly tool performance we constructed poreTally, a comprehensive benchmarking tool. poreTally automatically assembles a given read set using several often-used assembly pipelines, analyzes the resulting assemblies for correctness and continuity, and finally generates a quality report. Results can immediately be shared with peers in a Github/Gitlab repository. Furthermore, we aim to give a more inclusive overview of assembly pipeline performance than any individual research group can, by offering users the possibility to submit their results to a collective benchmarking effort. poreTally is available on Github.

## 1 Introduction

Nanopore sequencing is a third-generation nucleic acid sequencing method that produces error-prone long reads of consistent quality. In short, a DNA or RNA strand is pulled through a protein pore while the characteristic way in which the bases influence the electrical resistance through the pore is recorded. Base callers, software tools that use empirically obtained information on the characteristic current levels for certain combinations of bases, can then be used to translate the electric current over time into a nucleic acid sequence.

Nanopore sequencing was first brought to commercial practice in 2014, with the introduction of the handheld MinION sequencer by Oxford Nanopore Technologies (ONT). Two upscaled platforms, GridION and PromethION, followed soon after. Users subsequently developed various open-source analysis tools that were attuned to the greater read length and high sequencing error rates that characterized nanopore reads initially (approximately 75% [17]). As was to be expected of the first attempt at a radically different approach to sequencing, the sequencing hardware, base callers and sample preparation practices underwent many changes since their introduction. As a cumulative result of all improvements so far, unprocessed MinION reads have now been reported to reach an identity rate of 93% in the hands of users [11], with a length distribution around 10 kbase [18], although these characteristics have been known to vary with sample preparation method and species [20, 21].

Although ONT’s update schedule provides the user community with tools of steadily increasing value, it requires downstream analysis tool developers to repeatedly re-parameterize their tools. Providing updated benchmarks for said tools has proven difficult as well. In the case of *de novo* assembly pipelines, several benchmarks of the most popular tools at a given point in time have been run and published in peer-reviewed journals [9, 4, 6, 1]. However, in each case ONT significantly improved some aspect of read quality before or shortly after the publication appeared, rendering it partially outdated. Furthermore, each benchmarking effort focused on one organism, while it has been shown that read quality and best assembly practices may differ from one taxon to the next [19, 15, 16]. As ONT intends to continue frequent improvement of its platforms, no single research group can realistically be expected to benchmark *de novo* assembly pipelines for a reasonably diverse set of organisms and publish their work in traditional academic journals, without the next major update to ONT’s hardware and software being close.

To keep up with the rapid pace of developments in nanopore sequencing, the ONT user community has broadly adopted several strategies in disseminating results and new tools. For example, the community embraced publication on preprint servers such as BioRxiv, while publication in traditional peer-reviewed journals is rather done to formally confirm validity of the work long after the user community has done its own informal evaluation [e.g. 7, 8]. Similarly, the fast open peer-review system offered by publisher F1000 proved moderately popular, as evident from the activity on their publication channel dedicated to nanopore sequencing. Recently, Wick *et et al.* took an interesting approach to share their findings on MinION base caller performance, by publishing their analysis on Github and regularly updating their benchmark as new developments unfolded [22]. A more collaborative effort was adopted for nucleotid.es, a benchmarking platform for short read assembly pipelines that allows users to submit Docker images of pipelines and evaluates these on standard datasets. nucleotid.es thus facilitates iterative benchmarking and an objective comparison of pipelines. However, it does not allow benchmarking on long error-prone reads or user-supplied datasets, nor does it support essential practices for nanopore assemblies, such as post-assembly polishing using raw nanopore signal.

To address both the rapid devaluation of *de novo* assembly pipeline benchmarks due to hard- and software updates and the variability of their performance across taxa, we propose a community-driven frequent benchmarking practice and present a tool to facilitate this. We encourage research groups that make use of nanopore sequencing to benchmark assembly pipelines for their organisms of interest following a standardized routine and publish their results directly online. This will provide other users working on the same or similar taxa with an indication of the best assembly pipeline for their case with the most up-to-date hard- and software. Our tool, poreTally, supports this practice by offering often-used assembly pipelines, a performance analysis routine, report generation and publication on free repository hosting services Github or Gitlab in one package, providing total transparency throughout the process. poreTally was designed to allow easy extension to other pipelines or multiple instances of the same pipeline with different parameter settings. Optionally, users may submit their results to a collective benchmark effort. Submitted benchmark results will periodically be summarized and presented to the community as a measure of overall performance and a choice guide for particular user cases. Of course, submitters will be duly credited and referred to.

## 2 Methods

poreTally is written in Python 3 (Python Software Foundation, https://www.python.org/). Its workflow is divided in three steps. First, it executes a number of assembly pipelines on a provided dataset. Then several analysis tools are run on the produced assemblies. In the final step, summarized results are automatically published on Github or Gitlab in a readable format. A schematic overview of the process is given in Figure 1.

**Figure 1:**
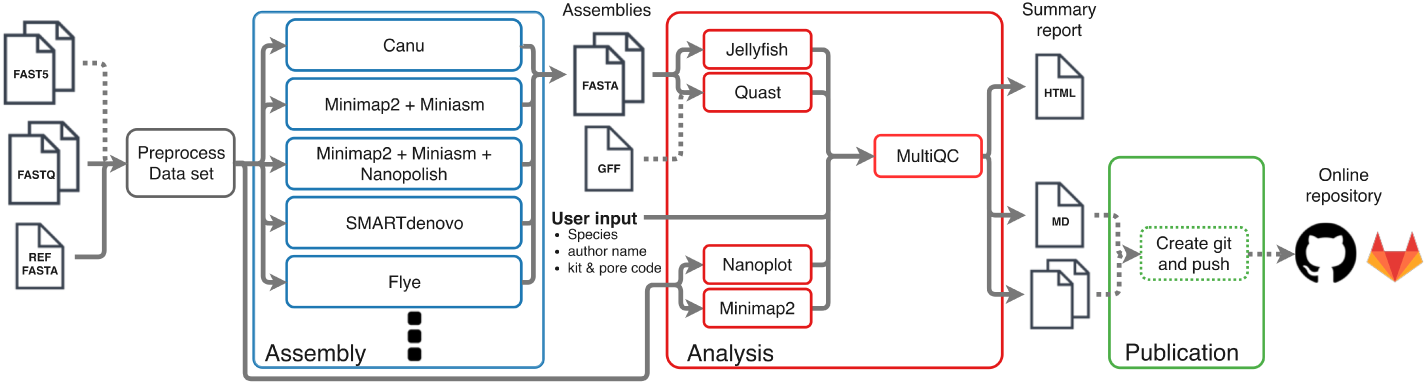
Schematic representation of the poreTally workflow. Optional steps are denoted with dotted lines. poreTally requires FASTQ-files of Nanopore reads and a reference genome. After joining reads in multi-FASTQ’s and determining genome coverage using the reference genome, reads are fed in parallel to several assembly pipelines. The resulting assemblies are analyzed using Jellyfish (version 2.2.7) [14] and Quast (Version 4.6.3) [5]. Optionally, a reference gene annotation file (GFF) may be supplied to determine how many known genes could be retrieved from the assemblies. The quality of the read set is analyzed using Nanoplot [2] and aligned to the reference genome using Minimap2 (version 2.9-r720) [12]. The generated analysis reports are summarized by MultiQC [3] and can be shared on Github/Gitlab.

### 2.1 Installation

poreTally can be run using the provided docker container or by installing it through pip. In the latter case, some minimal requirements have to be met first; Python 3.6, a miniconda/anaconda installation and Git.

### 2.2 Assembly pipeline preparation and running

For the running of assembly pipelines, poreTally relies on the Snakemake workflow management system [10] and its excellent integration with conda environments. The user is required to provide at least a reference genome and nanopore reads in FASTQ-format. The original FAST5-format reads must be supplied as well, should some tools in the pipelines require this (e.g. Nanopolish [13]). Genome size and coverage are automatically derived from the input files for tools that require these parameters. Next, a Snakemake workflow is generated containing one independently executable set of commands for every assembly pipeline. If required, dedicated conda environments containing the necessary tools for a particular pipeline are generated on the fly. Pipelines are then run while memory usage and wall time are monitored. By default, pipelines are run sequentially, but if a SLURM scheduler is installed on the system pipelines can be run in parallel as separate SLURM jobs by providing a JSON-file containing the required header information. poreTally is pre-configured to run several often-used assembly pipelines, however users can easily run other tools by providing YAML-files containing the commands that should be run and build information for a conda environment if required.

### 2.3 Pipeline performance analysis

After the assembly pipelines have finished running, the quality of the original read set and of the produced assemblies is assessed. Similarly to assembly pipelines, analyses are executed as a Snakemake workflow and can be parallelized using SLURM.

Nanoplot [2] is used to assess raw read length and quality scores. Minimap2 [12] then maps the reads to the reference genome to estimate deletion, insertion and substitution error rates. For the assessment of assembly quality, poreTally relies on a combination of Quast [5] for assembly continuity measures (e.g. N50) and error rates, and Jellyfish [14] for five-mer frequency counts, which may elucidate a bias towards certain sequences. Finally, results are summarized in an interactive HTML-report using MultiQC [3] (a custom fork of version 1.5.dev0, provided together with poreTally) with an adapted report template and several custom modules (Figure 2). As the HTML-report does not display properly if results are uploaded to Github or Gitlab, a static markdown version is also generated.

**Figure 2:**
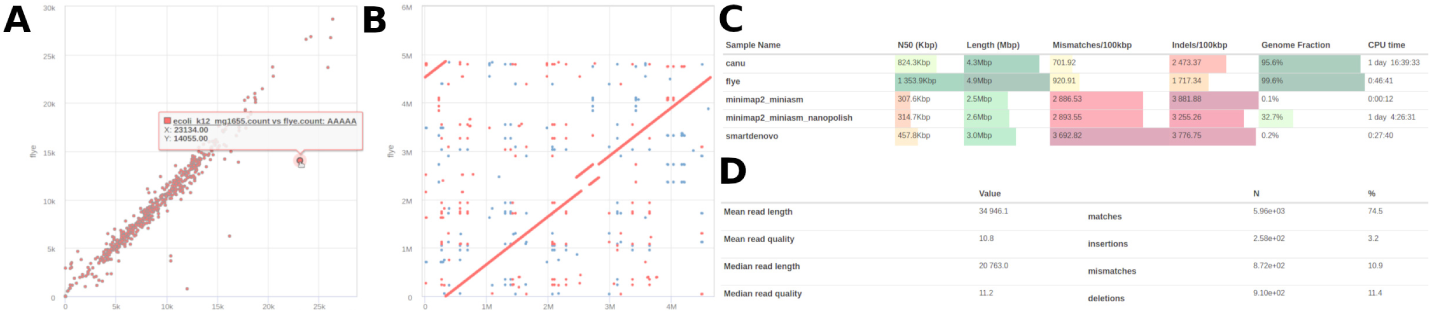
Examples of four elements in the report produced by poreTally: a *k*-mer abundance graph **(B)**, synteny plot **(C)**, assembly pipeline performance metrics **(D)** and read set quality measures **(E)**.

### 2.4 Publication

If a git repository address was provided, the results are now uploaded to Github or Gitlab. If the user chooses to allow the use of their results in the collective benchmarking effort, a specific empty repository will be forked and the generated results will be submitted as a new pull request to this repository. Periodically, the made pull requests will be accepted, after which we will summarize and visualize the results, and present them in a separate Github repository.

## 3 Discussion

ONT’s nanopore sequencers are a novelty in nucleic acid sequence analysis in many respects, but while the long reads, low initial investment cost and small physical footprint are praised frequently, the unusual development of its user community is mentioned less often. First, the community was actively involved in the development of nanopore sequencing platforms. As a result, many of its members have the required knowledge of, and access to the hardware and software to modify every step in the sequencing process. Second, the user community has adapted to the rapid development of their field by embracing faster alternatives to traditional peer-reviewed journals as its main means for publication.

In line with these developments, we present poreTally to provide the user community with a faster, more flexible alternative to peer-reviewed benchmark papers for *de novo* assembly tool performance. poreTally is an easy to use framework which aims to standardize the entire benchmarking process, by implementing *de novo* assembly pipelines, assembly evaluation and report publication in one package. By lowering the effort required for each of these steps, we aim to motivate evaluation and frequent re-evaluation of assembly pipelines, and publication of independent benchmarks. As the number of published reports increases, the probability of finding one that pertains to a particular user’s case, i.e. on a related organism, with current hardware and software versions, should increase as well.

A potential downside of the publication process as it is currently implemented, is that it may give a fragmented view of assembly pipeline performance. Therefore, we also included the option to submit results to a collective benchmark effort. Results submitted to this effort will be summarized, thus giving a more unified indication of *de novo* assembly pipeline performance.

In conclusion, poreTally enables the nanopore user community to collectively map the landscape of available *de novo* assembly pipelines, thus providing a choice guide for a wide array of user cases. As the per-base cost of nanopore sequencing continues to decrease and the potential to sequence new species grows, we expect the importance of such guidelines to grow as well.

## References

[1] D. S. Cali, J. S. Kim, S. Ghose, C. Alkan, and O. Mutlu. Nanopore sequencing technology and tools for genome assembly: computational analysis of the current state, bottlenecks and future directions. Briefings in Bioinformatics, 2018. doi:10.1093/bib/bby017. on-line pre-publication, accessed 2018-04-04.

[2] W. De Coster, S. D’Hert, D. T. Schultz, M. Cruts, and C. Van Broeckhoven. NanoPack: visualizing and processing long read sequencing data. bioRxiv, 2018. doi:10.1101/237180.

[3] P. Ewels, M. Magnusson, S. Lundin, and M. Käller. MultiQC: summarize analysis results for multiple tools and samples in a single report. Bioinformatics, 32(19): 3047–3048, 2016.

[4] F. Giordano, L. Aigrain, M. A. Quail, P. Coupland, J. K. Bonfield, R. M. Davies, G. Tischler, D. K. Jackson, T. M. Keane, J. Li, et al. *De novo* yeast genome assemblies from MinION, PacBio and MiSeq platforms. Scientific Reports, 7(1): 3935, 2017.

[5] A. Gurevich, V. Saveliev, N. Vyahhi, and G. Tesler. QUAST: quality assessment tool for genome assemblies. Bioinformatics, 29(8):1072–1075, 2013.

[6] B. Istace, A. Friedrich, L. d’Agata, S. Faye, E. Payen, O. Beluche, C. Caradec, S. Davidas, C. Cruaud, G. Liti, et al. *De novo* assembly and population genomic survey of natural yeast isolates with the Oxford Nanopore MinION sequencer. GigaScience, 6(2):1–13, 2017.

[7] M. Jain, S. Koren, J. Quick, A. C. Rand, T. A. Sasani, J. R. Tyson, A. D. Beggs, A. T. Dilthey, I. T. Fiddes, S. Malla, H. Marriott, K. H. Miga, T. Nieto, J. O’Grady, H. E. Olsen, B. S. Pedersen, A. Rhie, H. Richardson, A. Quinlan, T. P. Snutch, L. Tee, B. Paten, A. M. Phillippy, J. T. Simpson, N. J. Loman, and M. Loose. Nanopore sequencing and assembly of a human genome with ultra-long reads. bioRxiv, 2017. doi:10.1101/128835.

[8] M. Jain, S. Koren, K. H. Miga, J. Quick, A. C. Rand, T. A. Sasani, J. R. Tyson, A. D. Beggs, A. T. Dilthey, I. T. Fiddes, et al. Nanopore sequencing and assembly of a human genome with ultra-long reads. Nature Biotechnology, 36(4):338, 2018.

[9] K. Judge, M. Hunt, S. Reuter, A. Tracey, M. A. Quail, J. Parkhill, and S. J. Peacock. Comparison of bacterial genome assembly software for MinION data and their applicability to medical microbiology. Microbial Genomics, 2(9), 2016.

[10] J. Köster and S. Rahmann. Snakemake—a scalable bioinformatics workflow engine. Bioinformatics, 28(19):2520–2522, 2012.

[11] M. Leija-Salazar, F. J. Sedlazeck, K. Mokretar, S. Mullin, M. Toffoli, M. Athanasopoulou, A. Donald, R. Sharma, D. Hughes, A. H. Schapira, et al. Detection of GBA missense mutations and other variants using the Oxford Nanopore MinION. bioRxiv, 2018. doi:10.1101/288068.

[12] H. Li. Minimap2: versatile pairwise alignment for nucleotide sequences. arXiv, 1708, 2017.

[13] N. J. Loman, J. Quick, and J. T. Simpson. A complete bacterial genome assembled de novo using only nanopore sequencing data. Nature methods, 12(8):733, 2015.

[14] G. Marçais and C. Kingsford. A fast, lock-free approach for efficient parallel counting of occurrences of k-mers. Bioinformatics, 27(6):764–770, 2011.

[15] T. P. Michael, F. Jupe, F. Bemm, S. T. Motley, J. P. Sandoval, C. Lanz, O. Loudet, D. Weigel, and J. R. Ecker. High contiguity *Arabidopsis thaliana* genome assembly with a single nanopore flow cell. Nature communications, 9(1): 541, 2018.

[16] D. E. Miller, C. Staber, J. Zeitlinger, and R. S. Hawley. High-quality genome assemblies of 15 *Drosophila* species generated using Nanopore sequencing. bioRxiv, 2018. doi:10.1101/267393.

[17] J. Quick, A. R. Quinlan, and N. J. Loman. A reference bacterial genome dataset generated on the MinION portable single-molecule nanopore sequencer. Giga-Science, 3(1):22, 2014.

[18] M. Schalamun, D. Kainer, E. Beavan, R. Nagar, D. Eccles, J. Rathjen, R. Lanfear, and B. Schwessinger. A comprehensive toolkit to enable MinION long-read sequencing in any laboratory. bioRxiv, 2018. doi:10.1101/289579.

[19] M. Schmid, D. Frei, A. Patrignani, R. Schlapbach, J. E. Frey, M. N. Remus-Emsermann, and C. H. Ahrens. Pushing the limits of *de novo* genome assembly for complex prokaryotic genomes harboring very long, near identical repeats. bioRxiv, 2018. doi:10.1101/300186.

[20] M. H.-W. Schmidt, A. Vogel, A. K. Denton, B. Istace, A. Wormit, H. van de Geest, M. E. Bolger, S. Alseekh, J. Maß, C. Pfaff, et al. *De novo* assembly of a new *Solanum pennellii* accession using nanopore sequencing. The Plant Cell, 29 (10):2336–2348, 2017.

[21] E. A. Solares, M. Chakraborty, D. E. Miller, S. Kalsow, K. E. Hall, A. G. Perera, J. Emerson, and R. S. Hawley. Rapid low-cost assembly of the *Drosophila melanogaster* reference genome using low-coverage, long-read sequencing. bioRxiv, 2018. doi:10.1101/267401.

[22] R. R. Wick, L. M. Judd, and K. E. Holt. Comparison of Oxford Nanopore basecalling tools, 2018. https://github.com/rrwick/Basecalling-comparison, accessed 2018-04-04.

